# Peptidoglycan Mediates Loa22 and Toll-like Receptor 2 Interactions in Pathogenic *Leptospira*

**DOI:** 10.1101/520288

**Authors:** Shen-Hsing Hsu, Ming-Yang Chang, Yi-Ching Ko, Li-Feng Chou, Ya-Chung Tian, Cheng-Chieh Hung, Chih-Wei Yang

## Abstract

Leptospirosis is an overlooked zoonotic disease caused by pathogenic *Leptospira.* The kidney is the major organ infected by *Leptospira* which causes tubulointerstitial nephritis. *Leptospira* outer membrane components contain several virulence factors that play important roles in the pathogenesis of leptospirosis. Among them, OmpA-like protein Loa22 is essential for leptospiral virulence. However, the pathogenic mechanisms of tubulointerstitial nephritis involving this virulence factor are still unclear and need further investigation. In this study, pull-down assays suggested that Toll-like receptor 2 (TLR2) proteins interacted with Loa22 from *Leptospira* outer membrane extractions. Combination of Atomic force microscopy (AFM) and side-directed mutagenesis suggested that Loa22 exhibited high affinity for *Leptospira* peptidoglycan (LPGN) and the residues of Loa22 were involved in LPGN interaction. Mutation of two key residues within the OmpA-like domain of Loa22, Asp^122^ and Arg^143^, significantly attenuated their relative affinities for LPGN indicating that these two residues were responsible for LPGN binding. Thus Loa22 OmpA domain was responsible for interacting with LPGN and the two indicated residues may participate in binding to LPGN. Recombinant Loa22 (rLoa22) protein was further complexed with LPGN and incubated with HEK293-TLR2 cells to monitor inflammatory responses. Inflammatory responses were provoked by rLoa22-LPGN complexes, but not rLoa22 alone, involved *CXCL8/IL8, hCCL2/MCP-1,* and *hTNF-α* activation. Confocal microscopy further identified the co-localization of Loa22-LPGN complexes and TLR2 receptors on HEK293-TLR2 cell surface. The affinity between Loa22-LPGN complexes and TLR2 were further confirmed and measured by AFM and ELISA. Downstream signals from TLR2 including p38, ERK, and JNK were observed by western blotting induced by Loa22-LPGN complexes. In summary, this study identified LPGN in leptospira mediates interactions between Loa22 and TLR2 and induces downstream signals to trigger inflammatory responses. Interactions between Loa22-LPGN-TLR2 reveal a novel binding mechanism for the innate immune system and infection induced by leptospira.

**Author summary:** Leptospirosis is one of the most overlooked zoonotic diseases caused by pathogenic *Leptospira* in warm and humid regions worldwide. With the infection by *Leptospira,* many organs are invaded and can result in multiple-organ failure (Weil’s syndrome). Kidney is the major organ infected by pathogenic *Leptospira,* which would manifest as tubulointerstitial nephritis. In this study, we focused on the outer membrane lipoprotein Loa22 (*Leptospira*l OmpA-like domain 22) from pathogenic *Leptospira* which triggers inflammatory responses on renal tubular cell. Protein domain prediction indicated that Loa22 contains an important domain termed OmpA-like domain and the function of this domain is peptidoglycan (PGN) binding. From sequence alignments of Loa22 with other OmpA proteins, two important amino acids, Asp^122^ and Arg^143^, were found to be highly conserved. The role of the two conserved residues in AbOmpA (OmpA protein in *A. baumannii*) and Pal (peptidoglycan-associated lipoprotein in *E. coli)* proteins are important for PGN binding. These two residues in Loa22 were altered by site-directed mutagenesis to obtain D122A and R143A variants. In pull-down and AFM analysis, the binding capacities of Loa22 variants to *Leptospira* PGN (LPGN) were significantly decreased as compared to rLoa22WT, indicating that the two residues are involved in LPGN binding. Furthermore, recombinant Loa22 and its variants in the absence or presence of LPGN, were incubated with HEK293-TLR2 cells, to confirm the role of LPGN in triggering inflammatory responses involving *CXCL8/IL8, hCCL2/MCP-1,* and *hTNF-α.* These factors are involved in downstream signaling of inflammatory responses through Toll-like receptor 2 (TLR2). In addition, confocal microscopy was employed to observe the co-localization of Loa22-LPGN complexes and TLR2 receptors on HEK293-TLR2 cell surfaces. Finally, the interaction forces between rTLR2 and rLoa22-LPGN complexes were measured by AFM and ELISA to conclude the necessary role of LPGN in rLoa22-TLR2 complex formation. In summary, these results demonstrate that the interaction of Loa22 protein with the important cell wall component, PGN, concomitantly triggered inflammatory responses of host cells through interaction with TLR2.

## Introduction

*Leptospira* is the pathogen of the most overlooked zoonotic diseases leptospirosis, which results in multiple-organ failure (Weil’s syndrome), especially of the kidney (1, 2). The disease is generally transmitted through contact with urine of carrier hosts in water or soil, causing infection in humans via dermal or gastrointestinal routes (3). Clinical symptoms in humans include high fever, jaundice, and renal failure (4). The major targets of *Leptospires* in the kidney are the renal proximal tubular cells (2). One line of evidence has shown that pretreatment of kidney epithelial cells with outer membrane proteins from *Leptospira* triggers significant expression of tubulointerstitial nephritis-related genes (5). Surface-exposed antigens, due to their location, are likely involved in primary host-pathogen interactions, adhesion, and/or invasion (6). These interactions are followed by bacterial adhesion to tissues, immune responses, and eventually bacteria entering the host. The *Leptospiral* outer membrane contains antigenic components include lipoproteins, lipopolysaccharide (LPS), and peptidoglycans (PGN) (7). *Leptospira* surface components implicated in virulence include sphingomyelinases, serine proteases, zinc dependent proteases, collagenase (8), LPS (9), PGN (7), major outer membrane lipoprotein 32 (LipL32) (10), *Leptospira* immunoglobulin-like (Lig) proteins (11), *Leptospira* surface adhesion proteins (Lsa) (12), and Loa22 *(Leptospira* OmpA-like lipoprotein) (13).

Recently, the identification of immunogenic outer membrane proteins of pathogenic *Leptospira* has become an important topic of *Leptospira* research. Among these proteins, Loa22 was detected in pathogenic *Leptospires* but not in non-pathogenic *Leptospires,* indicating the probable involvement of this protein in virulence (14). Previous studies showed Loa22 protein exhibits a bipartite structure, including an N-terminal domain (residues 1-77) and an OmpA domain (residues 78–186; predicted as peptidoglycan-associating motif) (Figure 1A). According to SpLip predictions, Loa22 is a lipoprotein, with a lipid modified Cys^21^ residue, and a cleavage site between residues 20 and 21 to form a mature lipoprotein (15, 16). The lipopeptide moieties of spirochetes are potent mediators of inflammatory responses. The Loa22 lipoprotein could contribute to innate immunity and may therefore induce severe disease manifestations by eliciting host immunopathogenic responses (17). Recombinant Loa22 was shown to bind to the extracellular matrix (ECM) including plasma fibronectin and collagen types I and IV *in vitro,* suggesting that the surface-exposed domain of Loa22 may act as an adhesin (12). The role of Loa22 during pathogenesis remains to be determined and the biological function and detailed mechanism involved in infection of host cells by *Leptospira* need further investigation.

**Figure 1.**
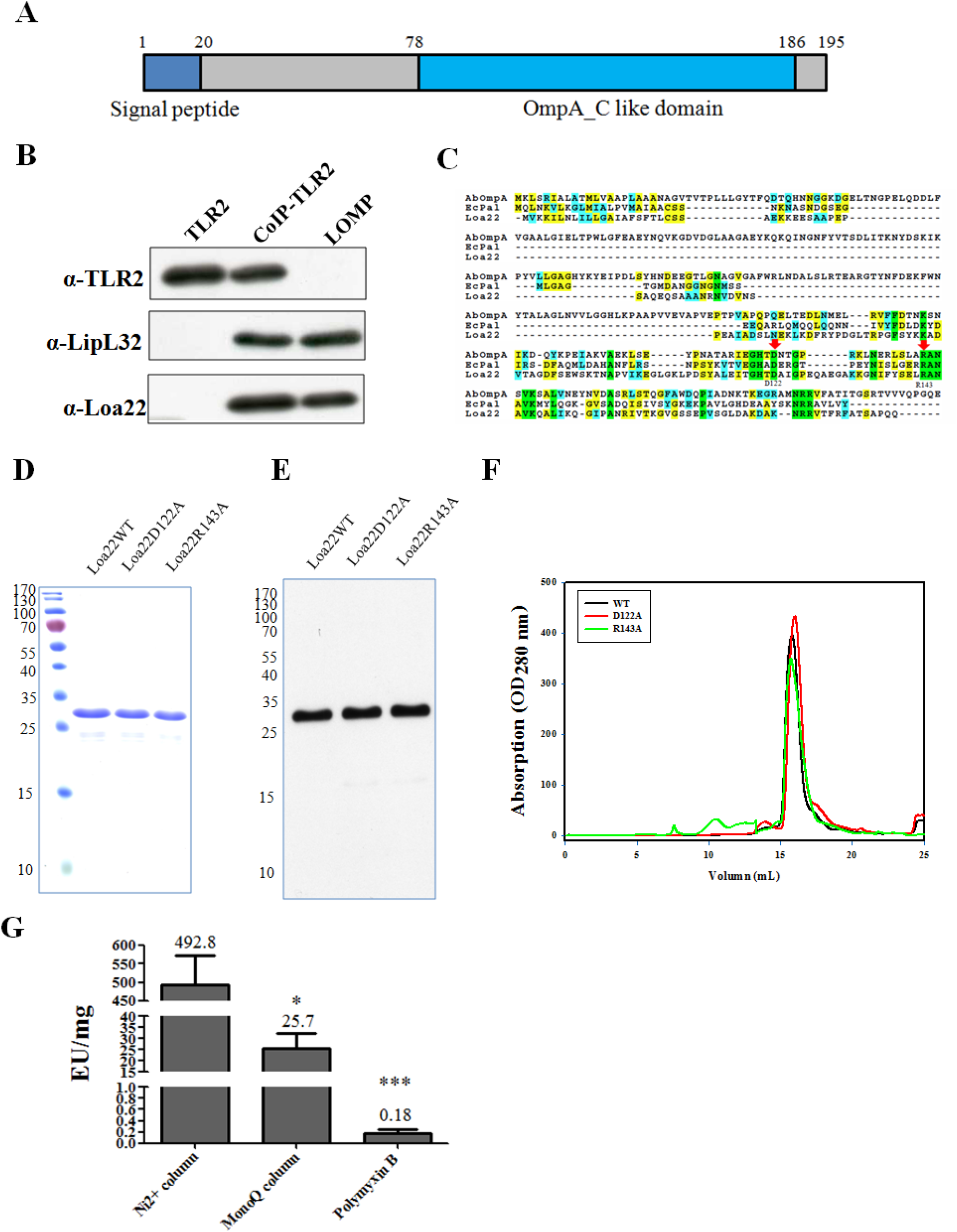
Characterization and bioinformatic analysis of Loa22 protein. (A) Domains prediction of Loa22. Loa22 contains 195 amino acids and the domains preditions revealed that the N terminal contains signal peptide (1–20) and C terminal of contains OmpA_C like domain (78-186). (B) Immuno-precipitant of purified TLR2 protein and Loa22 protein. Anti-TLR2, anti-LipL32 and anti-Loa22 antibodies were used to detect the presence of these proteins. (C) Sequence alignment of Loa22. Loa22 sequence was aligned with other OmpA domain protein including ABOmpA and Pal protein from *A. baumannii* and *E. coli,* respectively. Two pivotal residues (Asp122 and Arg143; red arrows) were highly conserved in Loa22 and other OmpA domain proteins and the two residues were responsible for peptidylglycan (PGN) binding. (D) Purification of the Loa22 and mutation variants. The molecular mass of Loa22 protein was calculated as 22 kDa and the SDS-PAGE showed the single band slightly higher than 25 kDa. (E) Western blot of purified Loa22WT and variants. The Loa22 WT and mutation variants were recognized by anti 6XHis tag antibody. (F) Size exclusion chromatography of purified Loa22 protein. The single peak of purified Loa22 protein with molecular mass about 22 kDa indicated that the protein is a uniform conformation. (G) LAL test of the purified recombinant Loa22. The endotoxin of the recombinant protein was assay by LAL test to measure the endotoxin contamination.* p<0.05; *, p<0.01;, p<0.001.

The initial interactions between pathogens and host cells trigger innate immune responses at the infection site (innate immunity is the first line of defense against bacterial infection in vertebrates) (18). Innate immune system develops germline-encoded pattern-recognition receptors (PRRs) to sense virulence components derived from various microbes (1). PRRs are responsible for recognizing microbe-specific molecules known as pathogen-associated molecular patterns (PAMPs) (19). Among the innate immunity systems, Toll-like receptors (TLRs) have been well studied in order to identify their function as PRRs (19). In humans, there are 10 TLR members (TLR1–TLR10) and 12 (TLR1–TLR9, TLR11–TLR13) in mouse. TLRs are further divided into two subfamilies; cell surface TLRs and intracellular TLRs, according to their localization. Cell surface TLRs include TLR1, TLR2, TLR4, TLR5, TLR6, and TLR10, whereas intracellular TLRs include TLR3, TLR7, TLR8, TLR9, TLR11, TLR12, and TLR13 (18). TLR family proteins play a pivotal role in innate immunity by recognizing conserved patterns in diverse microbial molecules (20). Among these TLRs, TLR2 in association with TLR1 or TLR6 is essential for sensing bacterial lipoproteins and lipopeptides (21, 22). TLRs containing leucine-rich repeats (LRRs) are responsible for pattern recognition at the extracellular portion and a Toll/IL-1 receptor (TIR) domain is responsible for signal transduction at the cytoplasmic portion (18). The crystal structure of TLR2/1 complex with its ligand, Pam3CSK4, revealed that two fatty esters of the glycerol moiety are embedded in a hydrophobic pocket of TLR2, while the amide-bound lipid chain is fitted into the hydrophobic channel in TLR1 (23). A lipid molecule of the lipoprotein presumably supported the binding domain of TLR2. In addition to the lipidation domain of lipoprotein, several known structures and motifs of the TLR2-binding protein have been reported. The PorB protein from *N. meningitides* has been suggested as a TLR2 ligand and the binding mechanism was hypothesized to involve electrostatic interactions contributing to ligand/receptor interactions (24). The BspA protein from *T. forsythia* with a LRR domain was also suggested as the TLR2 a ligand (25). The pentameric B subunit of type IIb *E. coli* enterotoxin (LT-IIb-B5) uses its hydrophobic upper pore region to directly bind TLR2 and the ring with four residues (Met^69^, Ala^70^, Leu^73^, and Ser^74^) defined as TLR2-binding sites for induction of inflammation responses (26). The *Leptospira* outer membrane lipoprotein, LipL32, interacts with TLR2 through Nβ1β2 and Cα4 domains and Val^35^, Leu^36^ and Leu^263^ were involved in TLR2 interaction (27). These TLR2 ligands use various binding mechanisms to interact with the innate immune system inducing inflammatory responses.

In *Leptospira,* it has been proven that Loa22 stimulates inflammatory responses and deletion of Loa22 from pathogenic *Leptospira* attenuate normal toxicity, while re-expression of the protein in *Leptospira* restores the *Leptospiral* virulence (13). However, the TLR2 binding domains and binding mechanisms of Loa22 are still unclear. In this study, pull-down assays revealed that the TLR2 protein interacted with Loa22 in the *Leptospira* outer membrane. Loa22 protein was also observed to co-localize with TLR2 on HEK293 cells over-expressing TLR2 at the surface, as detected by confocal microscopy. The corresponding inflammatory responses provoked by Loa22 were measured by real-time PCR. The Loa22 protein was further used to investigate interaction domains involving LPGN. We further mutated the two key OmpA domain residues, Asp^122^ and Arg^143^, and their relative affinities to LPGN. The OmpA domain was responsible for LPGN interactions and the two vital residues might participate in LPGN binding. The interactions of Loa22 and TLR2 were explored by ELISA and AFM, and the role of LPGN was verified to identify the interaction between Loa22 and TLR2. The binding mechanism of Loa22 and TLR2 is still unclear; therefore, we hypothesized that the OmpA domain of Loa22 plays vital roles in interactions with TLR2. It has been reported that TLR2 interacts with PGN molecules in inducing the inflammatory responses (28). Therefore, Loa22 was proposed to interact with LPGN molecules through the vital residues, and consequently interact with TLR2 to induce downstream signaling and cytokine production.

## Results

### Identification of TLR2 binding candidates from pathogenic *Leptospira*

Upon infection by pathogens, innate immune responses are induced to defense against the infection on host cell surfaces. We attempted to identify the TLR2 ligand from pathogenic *L. santarosai* serovar Shermani and characterize the binding mechanisms of virulence factors with TLR2. The human *TLR2* gene was sub-cloned from plasmid pUNO-TLR2 (Invivogen, San Diego, CA) and inserted into a lentivirus expression vector with a V5 tag at the C-terminus. The packaged virus particles were used to infect to HEK-293T cells, and a stable clone was selected using blasticidin for over-expression of the full-length human TLR2 protein. HEK-293T-TLR2 cells were used for full-length human TLR2 protein expression, and protein A-immobilized anti-V5 antibody was used for human TLR2 protein pull-down assays. After incubation of *Leptospira* outer membrane extractions with HEK-293T-TLR2 cells for two hours, the cells were lysed with lysis buffer, and protein A-immobilized anti-V5 antibody was used to pull-down the TLR2 and binding candidates. The pull-down fractions were analyzed by SDS-PAGE, and bands were extracted for MALDI-TOF analysis and protein identification. Several TLR2 binding candidates were isolated and one of the positive controls, LipL32, was observed by the proteins identification and anti-LipL32 recognition (Figure 1B) (27, 29, 30). This result demonstrated that the method used for identification of TLR2-binding candidates searching and identification is suitable and valid. An interesting TLR2 binding candidate, Loa22, was observed in the MALDI-TOF analysis and protein identification after co-immunoprecipitanting with TLR2. In order to confirm the MALDI-TOF results from co-immunoprecipitantition, an anti-Loa22 antibody was used to verify the interaction of Loa22 and TLR2. The western blot clearly demonstrated the interaction of Loa22 and TLR2 after co-immunoprecipitantion (Figure 1B). Loa22 was present in pathogenic *Leptospires* but not in non-pathogenic *Leptospires,* indicating that Loa22 protein is probably a virulence factor (14). Loa22 is anchored to the outer membrane of pathogenic *Leptospires* and contains a large OmpA domain, known as a peptidoglycan-binding domain (Figure 1A). Therefore, recombinant Loa22 (rLoa22) was constructed and expressed in *E. coli* to obtain purified rLoa22.

### Protein Purification and Mutagenesis

The Loa22 protein contains 195 amino acids, and domain prediction indicated the N-terminal signal peptide and C-terminal OmpA domain (Figure 1A). Sequence alignments of Loa22 with other OmpA domain proteins (Pal protein from *E. coli* and OmpA protein from *A. baumannii*) indicated that sequence similarity is low; however, two important PGN-binding residues, Asp^122^ and Arg^143^, were highly conserved in these OmpA domain proteins (Figure 1C). Therefore, these two residues were mutated to Ala using site-directed mutagenesis to generate D122A and R143A variants. The rLoa22 protein was expressed in *E. coli* ClearColi^TM^BL21 (DE3) pLys (Lucigen, Middleton, WI) and further purified by Ni^2+^-NTA affinity column and size exclusion chromatography (Figure 1D-F). In order to remove *E. coli* endotoxin contamination, the MonoQ column and polymyxin B resin were used to further remove endotoxin from purified rLoa22 protein. In the Limulus amebocyte lysate (LAL) assay, rLoa22 from *E. coli* Clearcoli™ contained negligible endotoxin and was suitable for inflammation assays (Figure 2G) (27). In addition, the N-terminal His_6_-tag tail was removed by Enterokinase and size exclusion chromatography, according to the previous report (27), in order to observe the obvious effects of rLoa22 protein without the His tag. In addition, rLoa22 protein and its related variants were treated at 100 ^°^C for 30 min or digested by proteinease K and served as a negative control under several conditions.

### PGN binding assay

Loa22 is a lipoprotein with a C-terminal OmpA domain, which is speculated to bind the essential cell wall component, PGN. To verify the PGN-binding activity of rLoa22, AFM was used to investigate the interaction between rLoa22 and LPGN. The *Leptospira* was immobilized on a mica surface and washed three times with PBS buffer containing 0.1% (w/v) Triton X-114 to remove the outer membrane and expose the PGN layer (Figure S1A,B). The rLoa22-modified AFM tip was used to measure the affinity between rLoa22 and *Leptospira* cell wall. AFM force-distance curves were recorded to distinguish specific and non-specific interactions (Figure 2A). The specific interaction force-distance curves were selected to analyze interactions between rLoa22 and LPGN. In contrast, the tip only was used to measure the *Leptospiral* surface, as well as the rLoa22-modified AFM tip was used to measure affinity for the mica surface as a negative control. The interaction forces of the two controls were calculated as 26.3 ± 5.1 and 31.2 ± 4.7 pN, respectively (Supplemental S1C,D and Figure 2B). The interaction force between rLoa22WT and PGN from *Leptospira* was calculated as 58.2 ± 5.6 pN (Supplemental S1E and Figure 2B). In addition, the binding frequency between rLoa22WT and LPGN was calculated as 15.8% as compared to the mica surface as 2.1% (Figure 2B). This result clearly demonstrated the LPGN binding activity of rLoa22 from pathogenic *Leptospira.* In addition, the rLoa22 mutation variants were prepared on AFM tip for PGN binding activity measurement. As expected, the interaction forces and binding frequency of D122A and R143A variants displayed low basal levels of LPGN binding activity that indicated the two residues played vital roles in LPGN binding (Supplemental S1F-G and Figure 2B). The two rLoa22 variants exhibited gross impairment in LPGN binding ability as compared to rLoa22WT, suggesting their crucial roles in maintaining of LPGN binding by rLoa22. To investigate the PGN binding activity of rLoa22 to other different PGN molecules, commercially available PGN molecules including those from *E. coli* (EPGN), *S. aureus* (SPGN), and *B. subtilis* (BPGN) were selected for incubation with rLoa22 protein at 37 ^°^C for 30 min. In addition, the PGN molecule from pathogenic *L.santarosai* serovor Shermani (LPGN) was isolated as described in Materials and Methods to test its affinity for rLoa22 protein. After three centrifugation steps and washes, the pellets were subjected to SDS-PAGE and western blot analysis. The results indicated that rLoa22 showed high affinity for LPGN (Figure 2C). The mutated variants of rLoa22, D122A and R143A, showed low affinity for the four types of PGN molecules, indicating that the two residues are important for PGN binding. Taken together, these results demonstrated that PGN molecules from pathogenic *Leptospira* with high affinity for rLoa22, while other PGN molecules exhibited relatively low affinity for rLoa22 protein (Figure 2C). Therefore, LPGN was selected for subsequent studies.

**Figure 2.**
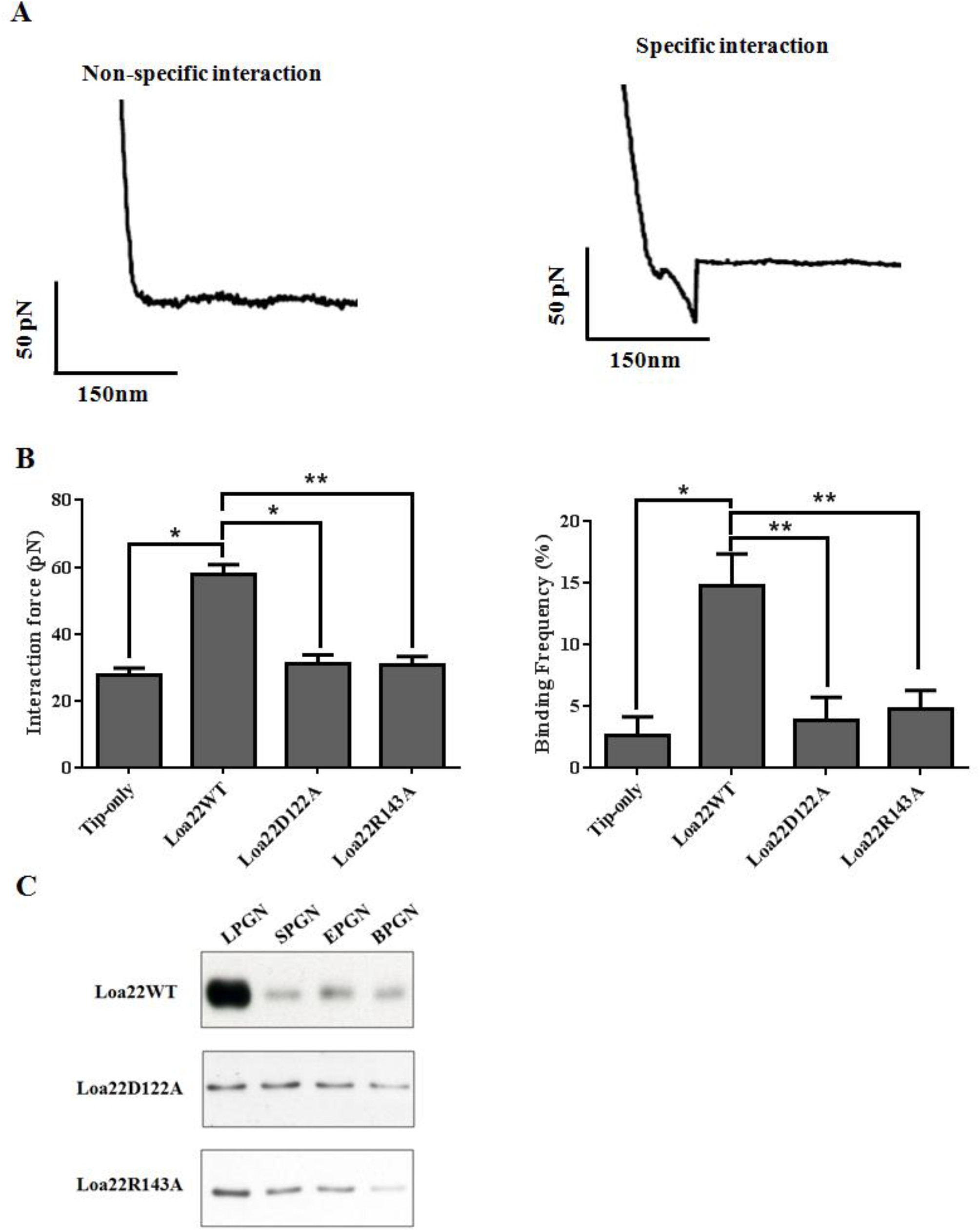
Rupture forces between rLoa22 and cell wall from *Leptospira* santarosai serovar Shermani surface as determined by smAFM. (A) The force-distance curves of AFM measurement. The tip-only (left panel, mock control) and rLoa22 (right panel, positive controls) modified AFM tip were used to analyze the interaction to *Leptospira* santarosai serovar Shermani cell wall. (B) The interaction forces and binding frequency of rLoa22 and *Leptospira* cell wall. rLoa22WT and mutation variants, D122A and R143A, were modified on AFM tip for rupture force measurements.(C) The PGN pull-down assay of rLoa22 variants and PGN molecules including the PGN from *Leptospira* santarosai serovar Shermani (LPGN), *E. coli* (EPGN), *S. aureus* (SPGN), and *B. subtilis* (BPGN), respectively. * p<0.05;, p<0.01;, p<0.001.

### rLoa22 Co-localizes with TLR2 on HEK293-TLR2 Cells

Attachment of *Leptospira* outer membrane proteins to host cell membrane is the first step invading the host during *Leptospira* infection. Previous studies showed that the Loa22 is up-regulation when host infection induces high levels of antibody production in the infected patient’s serum (14, 31). However, the receptor on host cell membrane which recognizes Loa22 protein is still unknown and needs further investigation. The results mentioned above suggested that TLR2 is a possible receptor on host cell membranes for Loa22. In order to demonstrate the co-localization of rLoa22 and TLR2, purified rLoa22 protein and its variants were incubated with HEK293-TLR2 cells for 4h and the cells were then washed, fixed, and incubated with conjugated antibodies for confocal microscopy analysis (Figure 3). rLoa22 and rTLR2 proteins were stained with rabbit polyclonal anti-Loa22 and mouse monoclonal anti-V5 primary antibodies follow by Alexa594 (red) conjugated anti-rabbit and Alexa488 (green) conjugated anti-mouse secondary antibodies, respectively. HEK293 cells lacking TLR2 expression were used as negative controls, with very little or no Alexa 488 fluorescence (Figure 3A). Co-localization of rLoa22WT-LPGN complexes and TLR2 receptors on HEK293-TLR2 cell was shown in Figure 3B. The results indicated that TLR2 receptor and the rLoa22 protein were mostly present on the cell surface, with consistent partial localization in the cytosol. The merged colors in several portions indicated that the two proteins were co-localized on HEK293-TLR2 cells (Figure 3B). In contrast, the two mutated variants reduced the cell binding ability, and the red color was absent in the confocal images (Figure 3C,D). The results from confocal microscopy clearly showed rLoa22-LPGN complexes directly interacted with TLR2 on HEK293-TLR2 cell surface, while the mutated variants, rLoa22D122A-LPGN and rLoa22R143A-LPGN, of rLoa22 significantly decreased co-localization with TLR2 on the cell surface.

**Figure 3.**
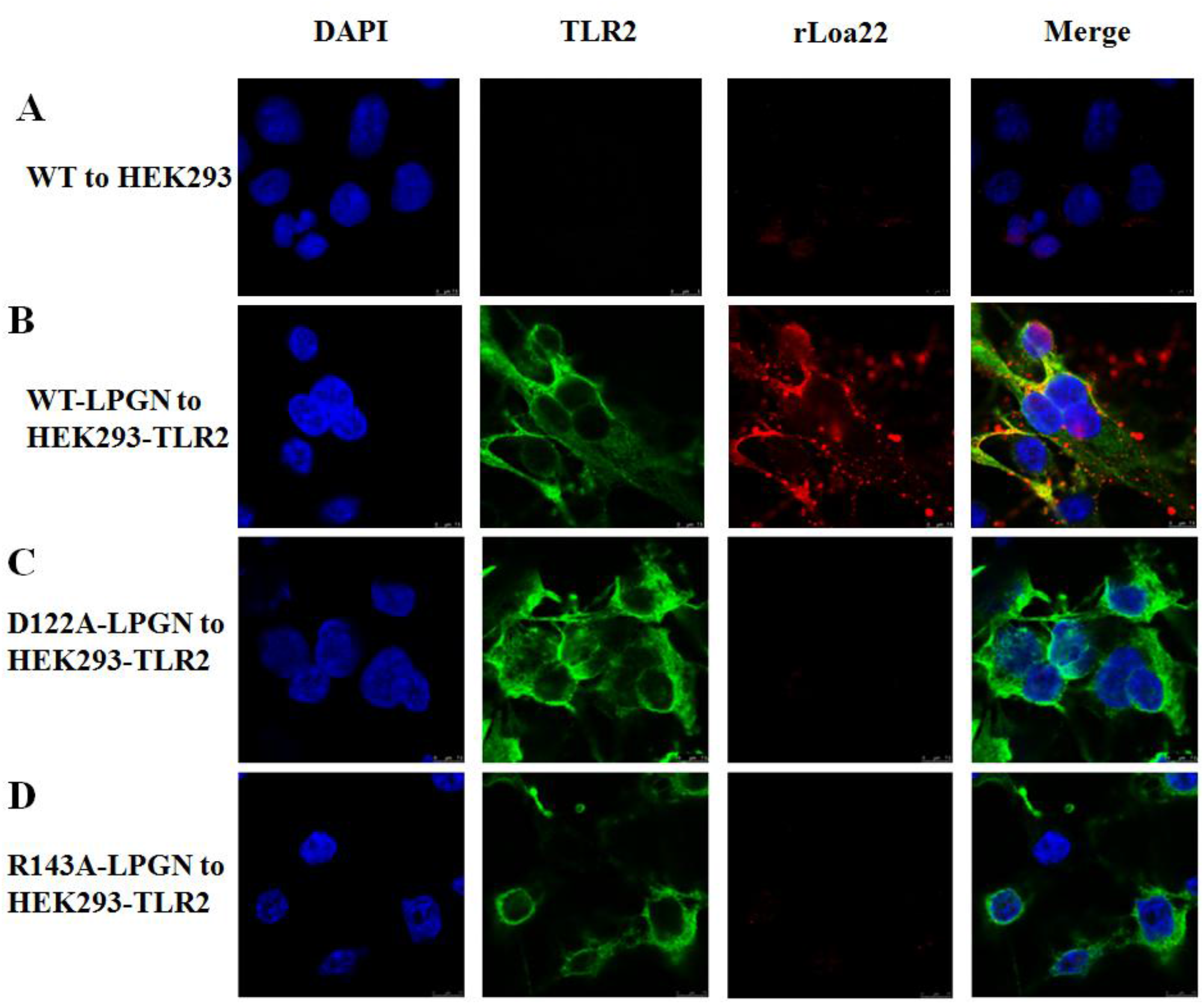
Loa22 co-localized with TLR2 on HEK293-TLR2 cells. The purified rLoa22 protein and its mutation variants with LPGN were incubated with HEK293-TLR2 cell for 4 h and the cell were fixed for confocal microscopy measurement. (A) rLoa22WT-LPGNwas incubated with HEK293 cell. (B) rLoa22WT-LPGNwas incubated with HEK293-TLR2 cell. (C) rLoa22D122A-LPGN was incubated with HEK293-TLR2 cell (D) rLoa22R143A-LPGN was incubated with HEK293-TLR2 cell. The nucleus was stained by DAPI (blue), TLR2 was stained by Alexa488 (green), and Loa22 was stained by Alexa594 (red). The yellow color indicated that the two proteins were co-localized in HEK293-TLR2 cell.

### Inflammatory Responses Induces by rLoa22-LPGN Complexes

Recognition of bacterial components by host TLRs initiates signaling cascades that stimulate nuclear transcription factor *κB* (NF-κB) and mitogen-activated protein kinases (MAPKs), and induces expression of chemokines and cytokines (29, 30). rLoa22 was expressed in *E. coli* ClearColi^TM^BL21 (DE3) pLys that contained low levels of endotoxin and was further purified by Ni^2+^-columns, MonoQ, and polymyxin to remove the contaminating endotoxin (Figure 2G). The expression levels of *CXCL8/IL8, hCCL2/MCP-1,* and *hTNF-α* were measured to investigate the role of rLoa22. Purified rLipL32 protein was used as the a positive control, and PBS buffer alone served as a negative control. Two hours after adding the rLoa22 to HEK293-TLR2 cells, the mRNA levels of *CXCL8/IL8, hCCL2/MCP-1,* and *hTNF-α* were significantly increased as compared with the PBS control (Figure 4A). rLoa22 stimulated the expression of *CXCL8/IL8, hCCL2/MCP-1,* and *hTNF-α* by 2.7-, 2.5-, and 4.1-fold, respectively. The results indicated that Loa22WT slightly increased the expression levels of *CXCL8/IL8, hCCL2/MCP-1,* and *hTNF-α.* LPGN was incubated with rLoa22 to form rLoa22-LPGN complexes and these were employed to stimulate HEK293-TLR2 cells. The rLoa22-LPGN complexes significantly increased expression levels of *CXCL8/IL8, hCCL2/MCP-1,* and *hTNF-α* as compared to PBS control, similar to the positive control LipL32 (Figure 4A) (27, 29). To exclude any His6-tag effect on Loa22, the His6-tag tail at the N-terminus was removed by Enterokinase, and the non-tagged rLoa22-LPGN complexes also showed significantly increased expression levels of *CXCL8/IL8, hCCL2/MCP-1,* and *hTNF-α* in HEK293-TLR2 cells as compared to PBS control. The results indicated that rLoa22-LPGN complexes, with or without the N-terminal His6 tag, showed similar behavior in inducing the expression of inflammatory cytokines. In order to prevent other effects of the His6-tag of rLoa22, non-His6-tagged proteins were used for subsequent studies. For further investigation of inflammatory cytokines induced by recombinant Loa22 protein, the rLoa22 was heat-treated (100 ^°^C, 30 min) and digested with proteinase K (20 μg/ml at 63 °C for 18 h), and the results revealed that there was no significant difference in stimulation of *CXCL8/IL8, hCCL2/MCP-1,* and *hTNF-α,* similar to the PBS control (Figure 4A). The absence of stimulatory effects after heat and proteinase K treatments further demonstrated that the rLoa22-LPGN complexes provoked the inflammatory responses.

**Figure 4.**
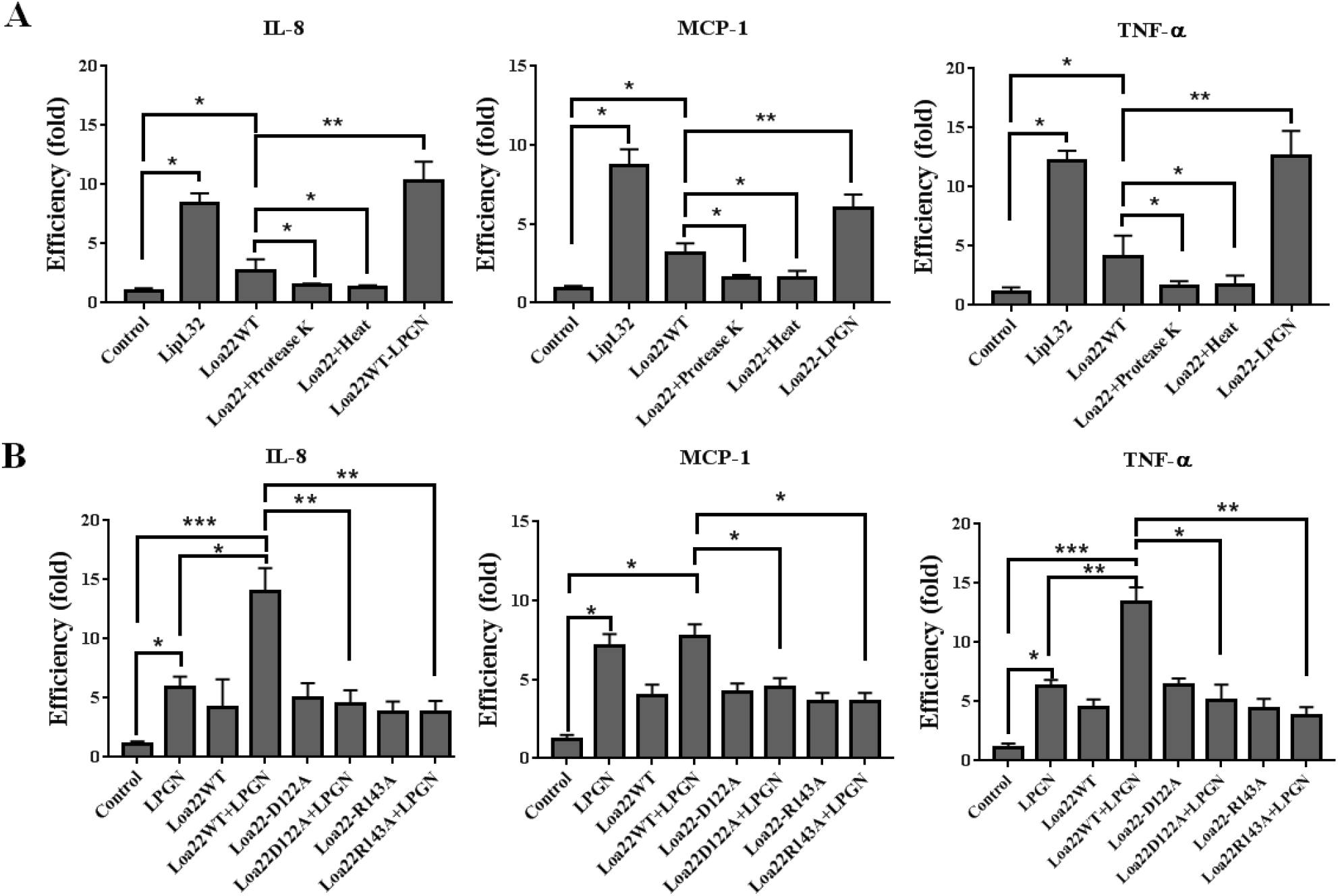
Inflammatory responses induced by rLoa22 protein in HEK293-TLR2 cell. The downstream inflammatory responses from TLR2 signaling such *CXCL8/IL8, hCCL2/MCP-1,* and *hTNF-α* were measured 2 hours after adding the stimulation agents. (A) Recombinant Loa22 isolated from Clearcoli^TM^ *E. coli* contained low level of endotoxin contamination and used for cytokines stimulation assay. LipL32 protein was used as positive control. The His6-Tag of rLoa22 was removed by Enterokinase protease. rLoa22 was treated with heat (100 ^°^C, 30 min) and proteinase K (20 μg/ml at 63 °C for 18 h) to denature and digested the protein. (B) Inflammatory responses stimulated by Loa22 and its variants in the presence or absence of PGN. rLoa22WT and mutated variants, rLoa22D122A and rLoa22R143A, were used to treat with HEK293-TLR2 cell in the presence or absence of LPGN.* p<0.05; *, p<0.01; *, p<0.001.

rLoa22 was the LPGN binding protein, and two vital residues were involved in PGN binding, as mentioned above. We further investigated the roles of LPGN and the mutated variants of rLoa22 in the stimulation of inflammatory responses. When treating HEK293-TLR2 cells with LPGN, expression of *CXCL8/IL8, hCCL2/MCP-1,* and *hTNF-α* were elevated by 5.9-, 7.1-, and 6.3-fold, respectively (Figure 4B). When HEK293-TLR2 cells were treated with rLoa22-LPGN complexes, the stimulation by *CXCL8/IL8, hCCL2/MCP-1,* and *hTNF-α* were significantly increased as compared to that in PBS controls (Figure 4B). The rLoa22-LPGN complexes significantly increased the expression levels of *CXCL8/IL8* and *hTNF-α* as compared to that in LPGN treatments, while the expression levels of *hCCL2/MCP-1* showed no significant differences between LPGN and rLoa22-LPGN complexes treatments (Figure 4B). These results indicated that the rLoa22-LPGN complexes stimulated high levels of *CXCL8/IL8, hCCL2/MCP-1,* and *hTNF-α* as compared to that of rLoa22 or LPGN alone. The rLoa22D122A and rLoa22R143A mutated variants with low affinity to LPGN showed significantly decreased expression levels of *CXCL8/IL8, hCCL2/MCP-1,* and *hTNF-α* as compared to that of rLoa22WT-LPGN complexes. The results further demonstrated that the LPGN cooperated with rLoa22 to interact with TLR2 and stimulated inflammatory responses.

### Interaction of TLR2 and rLoa22-LPGN complexes

Human TLR2 protein was expressed in HEK293-TLR2 cells and the protein was solubilized with 1% CHAPS in PBS buffer, and then purified by V5 antibody activated NHS-resin for affinity purification (Figure 5A). The purified TLR2 protein was used to bind the natural antagonist, Pam_3_CSK_4_, to measure binding activity. In addition, the major outer membrane lipoprotein, LipL32, was used to evaluate the binding activity of purified TLR2 protein (Figure 5B). In ELISA assays, the purified TLR2 protein showed high affinity for rLipL32 protein, but significantly decreased binding affinity for rLoa22. However, rLoa22-LPGN complexes significantly increased the affinity for TLR2, similar to rLipL32 protein (Figure 5B,C). In AFM measurements, the interaction force and binding frequency of rLoa22 to TLR2 were slightly increased as compared to BSA control (Figure 5D). However, the interaction force and binding frequency of rLoa22-LPGN complexes to TLR2 showed significantly increased as compared to BSA control (Figure 5D,E). These results provided direct evidence that rLoa22 cooperated with LPGN to interact with TLR2. In addition, the affinity of rLoa22 mutated variants, D122A and R143A, to TLR2 showed no significantly differences as compared to BSA control. It is not surprising that the rLoa22 mutation variants were loss of function variants that decreased the binding affinity for both LPGN and TLR2. Further addition of the LPGN molecules to these two variants could not raise the affinity to TLR2. By AFM measurements, the interaction forces and binding frequency of rLoa22 variants and TLR2 showed no significant differences as compared to rLoa22WT.

**Figure 5.**
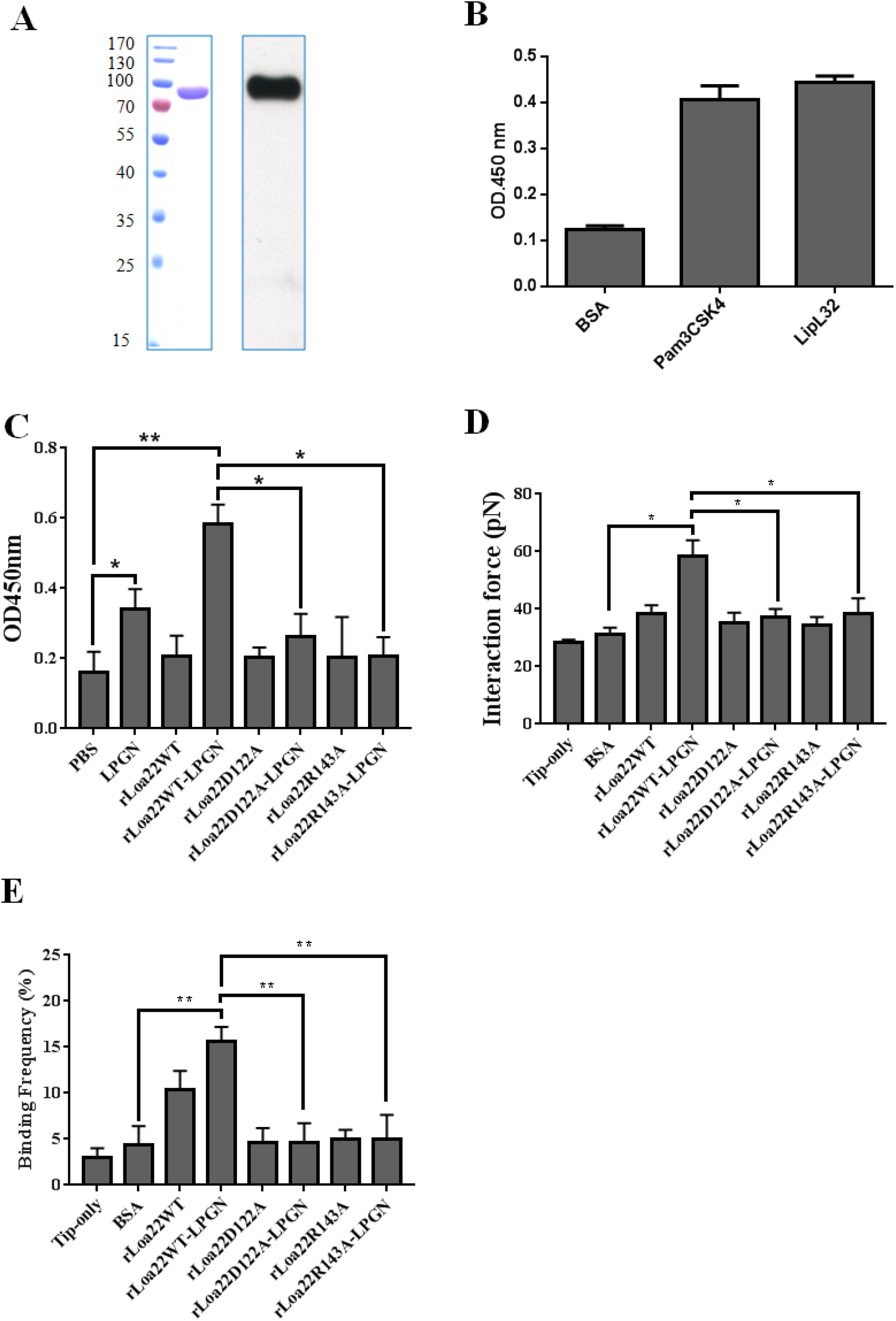
In vitro assay of the interaction between TLR2 and rLoa22-LPGN complexes. (A) Purification of TLR2 from HEK293-TLR2 cell. Left panel, SDS page of purified TLR2; right panel, western blot of purified TLR2. (B) Functional assays of purified TLR2 to Pam_3_CSK_4_ and *Leptospira* LipL32. (C) ELISA assay of the interaction between TLR2 and rLoa22. (D) AFM force-distance curves of the interaction between TLR2 and rLoa22. (E) The binding frequency of the interaction between TLR2 and rLoa22. TLR2 interacted to rLoa22WT and mutation variants, rLoa22D122A and rLoa22R143A in the presence or absence of LPGN. BSA was used as negative control and LipL32 was used as positive control.* p<0.05;, p<0.01; *, p<0.001.

### rLoa22-LPGN complexes induced p38, ERK, and JNK dependent signaling

*Leptospira* infection induces inflammatory responses through the TLR2-dependent pathway, and downstream signaling was therefore evaluated. The activation of the MAPK pathway was validated by western blot after treatment of HK2 and HEK293-TLR2 cells with rLoa22-LPGN complexes. The phosphorylation of MAPK pathway components including p38, ERK, and JNK was observed using their relevant antibodies. HK2 and HEK-293-TLR2 cells were cultured in serum free medium for 16 hours before adding the stimulating agent, rLoa22-LPGN complexes to precise evaluations of cellular function. Different time points were tested to determine the maximum phosphorylation levels of p38, ERK, and JNK. The maximum phosphorylation levels occurred at 1 hour after stimulation with rLoa22-LPGN complexes in serum-free HK2 and HEK-293-TLR2 cells. Stimulation of HEK293-TLR2 cells by rLoa22-LPGN complexes significantly increased the phosphorylation of p38, ERK, and JNK as compared to HEK-293 cells (Figure 6A). For TLR2 antibody neutralization experiments, HK2 cells were pretreated with anti-TLR2 antibody (10 μg/ml) for 1 hour, followed by adding of rLoa22-LPGN complexes for stimulation. Results in HK2 cells also revealed that cells pretreated with TLR2 antibody exhibited significantly decreased phosphorylation of p38, ERK, and JNK as compared to non-neutralized controls or non rLoa22-LPGN complexes stimulation controls (Figure 6B). All these results suggested that rLoa22-LPGN complexes stimulate the production of inflammatory responses through p38, ERK, and JNK signaling pathways.

**Figure 6.**
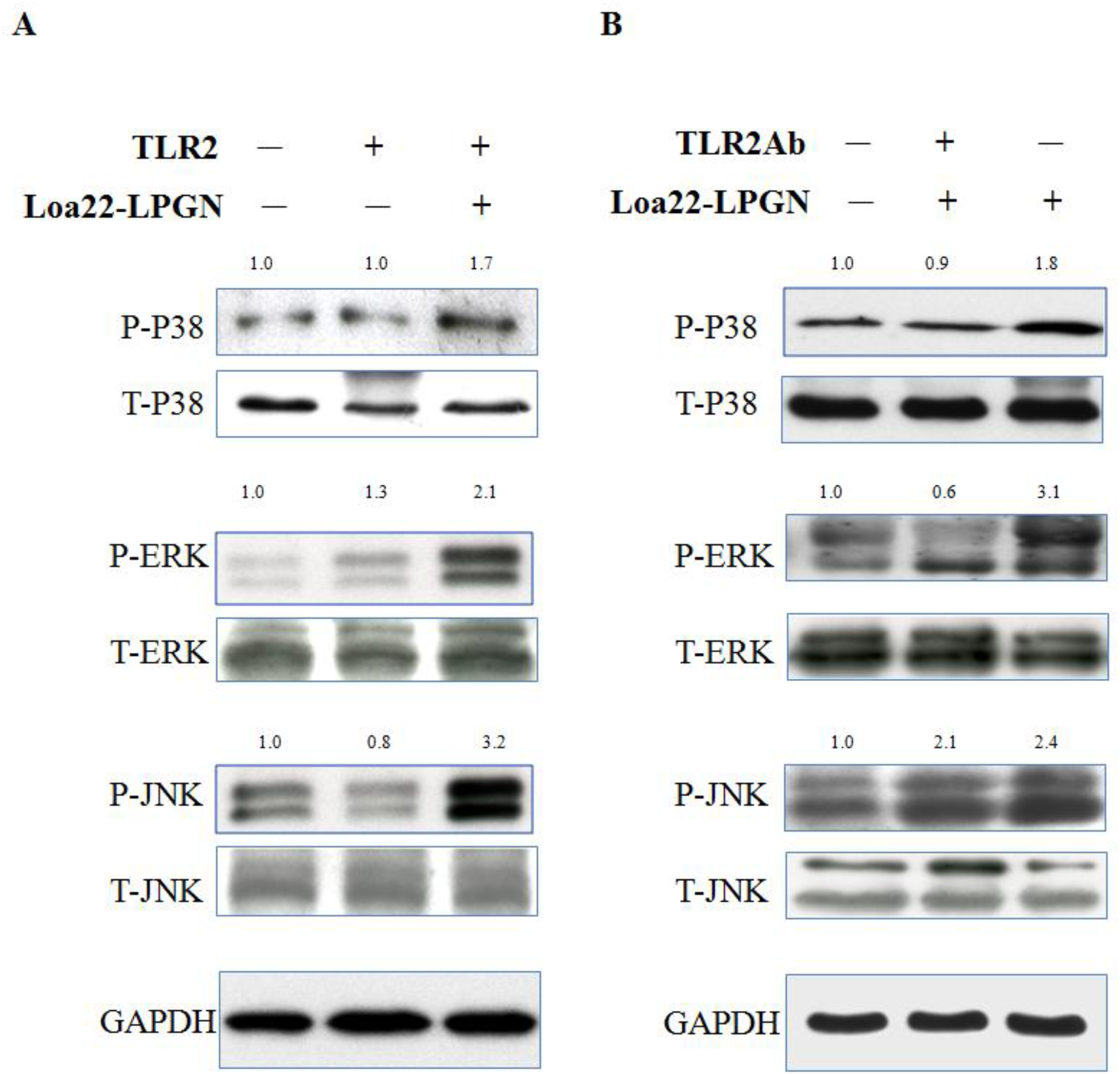
TLR2 downstream signaling cascades assays after rLoa22-LPGN complexes stimulation. (A) rLoa22-LPGN complexes stimulated HEK293 or HEK293-TLR2 cells for 24h and downstream signaling cascades were assayed. (B) rLoa22-LPGN complexes stimulated HK2 cells in the presence and absence of TLR2 antibody for 24h and downstream signaling cascades were assayed.

## Discussion

In *Leptospira,* Loa22 has been shown as an essential virulence factor, in that deletion of the Loa22 from pathogenic *Leptospira* attenuates the toxicity, whereas the re-expression the gene in *Leptospira* restores the virulence (13). However, the pathogenic mechanisms and the vital domains of Loa22 are still unclear. In this study, we used pull-down assays to demonstrate the interactions between TLR2 and Loa22 from *Leptospira* outer membrane extractions. One of the positive controls of this pull-down assay was LipL32, which has been proven as a TLR2 binding protein in pathogenic *Leptospira* (27, 29, 30). Interestingly, domain prediction of Loa22 protein suggested an OmpA-like domain and was speculated to interact with PGN. Therefore, PGN molecules were used to investigate the affinity of Loa22. However, PGN molecules from *E. coli* (EPGN), *S. aureus* (SPGN), and *B. subtilis* (BPGN) showed low affinity for Loa22. PGN molecules from pathogenic *Leptospira* (LPGN) are the only PGN molecules that showed relative high affinity for Loa22 (Figure 2C). These results suggested that the structure and composition of LPGN were different from those of EPGN, SPGN, and BPGN, and required further investigation. Combined with AFM force-distance curve studies and site-directed mutagenesis demonstrated that Loa22 directly interacted with LPGN on bacterial surfaces and two vital residues were involved in the interaction between Loa22 and LPGN (Figure 2). Furthermore, Loa22-LPGN complexes were observed to co-localize with TLR2 on HEK293-TLR2 cell surfaces as detected by confocal microscopy. The two mutated variants with low or no affinity for LPGN also revealed that the co-localization behaviors to TLR2 on HEK293-TLR2 cell surface was attenuated (Figure 3). In addition, the interaction forces and binding frequencies between rLoa22 and TLR2 showed no significant difference as compared to BSA control whereas the interaction forces and binding frequencies between rLoa22-LPGN complexes and TLR2 showed significant increases as compared to BSA control. The rLoa22 mutation variants also decreased the affinity for TLR2 in the absence or presence of LPGN as compared to rLoa22WT-LPGN complexes (Figure 5). We suggested two possibilities models concerning the relationships between rLoa22, LPGN, and TLR2. Firstly, Loa22 could not directly bind to TLR2; rather Loa22 used LPGN to interact with TLR2. According to this model, the concept of defining PGN as the TLR2 ligand is controversial, and whether PGN from pathogenic *Leptospira* as the TLR2 ligand is still unclear. The unique PGN from pathogenic *Leptospira* also showed higher affinities for Loa22 as compared to those from *E. coli, S. aureus,* and *B. subtilis* (Figure 2C). *Leptospira* is an idiographic bacterium that differs from other pathogens and its composition and reaction to host cells required further elucidation. Secondly, Loa22 interacted with LPGN and induced conformational changes in the protein, which exposed the TLR2-binding domain of Loa22 to interact with TLR2. The probability of the second model is higher in that the ligands may induce conformational changes in Loa22 and trigger the interaction between Loa22-LPGN and TLR2. Evidence for this model is that the Loa22 mutation variants exhibited low affinity for LPGN and a decrease of the affinity for TLR2.

The inflammatory responses provoked by rLoa22 in HEK293-TLR2 cells were used to measure the toxicity of this essential virulence factor. Previous studies have shown that *Leptospira* outer membrane proteins induce expression of nitric oxide, MCP-1, and TNF-α in cells (7). Tian *et al.* also demonstrated that *Leptospira* outer membrane proteins of *L. santarosai* Shermani increased collagen and TGF-β in renal proximal tubular cells (2). These data suggested that proinflammatory cytokine production might be involved in tubulointerstitial nephritis caused by *L. santarosai* Shermani infection through its outer membrane components. The recombinant protein method is the best way to investigate structural and functional relationships. However, endotoxin contamination from the recombinant protein can interfere with the intrinsic inflammatory properties of the innate immunity system. The endotoxin-free expression system provides the solution to overcome this problem. The ClearColi^TM^BL21(DE3) expression system was used to produce endotoxin free rLoa22 and its variants, and the yield of protein expressed was similar to that expressed in the BL21(DE3) expression system (32). In addition, anion-exchange chromatography (Mono-Q) and polymyxin B resin were further used to remove contaminating endotoxin from purified rLoa22 and its variants (Figure 1G). The LAL assay was used to confirm that the endotoxin was significantly removed. In terms of inflammatory responses, rLoa22WT protein showed low ability to induce inflammatory responses, whereas when rLoa22 combined with LPGN induced an increased inflammatory response, similar to the major outer membrane LipL32 protein (27). In fact, LPGN could significantly stimulate the expression of *CXCL8/IL8, hCCL2/MCP-1,* and *hTNF-α* (Figure 4B). Muller-Anstett *et al.* reported that SPGN co-localized with TLR2 and stimulated innate immune responses (28). The inflammatory responses induced by rLoa22WT-LPGN complex showed significantly increased as compared to PBS controls and LPGN (Figure 4). These results demonstrated that rLoa22-LPGN complexes effectively stimulated immune responses much more than LGPN alone, and further demonstrated that LPGN mediated the inflammatory responses induced by rLoa22. On the other hand, the rLoa22 mutation variants also reduced the ability to induce inflammatory responses in the absence or presence of LPGN as compared to rLoa22WT-LPGN complex stimulation (Figure 5). These results supported our hypothesis that LPGN bind rLoa22, further interact with TLR2 or LPGN bind to rLoa22, and induces conformational changes to interact with TLR2 on cell surfaces.

It has been reported that TLR2 interacts with PGN inducing the inflammatory responses (28). In this study, Loa22 was demonstrated to interact with LPGN through the two vital residues, and consequently interacted with TLR2 to induce downstream signal and cytokine production. Downstream signals induced by rLoa22-LPGN complexes were explored in HEK293 and HK2 cells, and MAPK pathway components, including p38, ERK, and JNK, were obviously stimulated. Previous studies have reported that HEK293 cell express no TLR2 on cell surfaces, and we constructed HEK293-TLR2 cells to express TLR2. HEK293 cells were used as normal controls. rLoa22-LPGN complexes significantly upregulated the phosphorylation levels of p38, ERK, and JNK in HEK293-TLR2 cells, but not in native HEK293 cells. In HK2 cells, adding anti-TLR2 antibody for neutralization procedure blocked the interaction between rLoa22-LPGN and TLR2, and therefore alleviated the activation of the MAPK pathway.

In summary, we demonstrated that Loa22 protein is a PGN binding protein. The PGN from *Leptospira* showed high affinity for Loa22. The LPGN binding activity of Loa22 is an important biological reaction to maintain cell wall stability and to protect against immune attack when infecting host cells. We further demonstrated that two key residues of the OmpA domain, Asp^122^ and Arg^143^, were involved in the affinity of Loa22 for LPGN. The interaction of Loa22 and TLR2 was explored by ELISA and AFM and the role of LPGN was verified to mediate Loa22 to interact with TLR2. Finally, Loa22 was proposed to interact with LPGN through two vital residues, and subsequently interacts with TLR2. This study identified LPGN in leptospira mediates interactions between Loa22 and TLR2 and induces downstream signals to trigger inflammatory responses. Interactions between Loa22-LPGN-TLR2 reveal a novel binding mechanism for the innate immune system and infection induced by leptospira.

## Materials and Methods

### Cell and Bacterial Culture

HK-2 (human kidney 2), immortalized human renal proximal tubular cell, was obtained from ATCC (number CRL-2190; Maryland, USA) and cultured in DMEM/Ham’s F12 (Life Technologies, Paisley, UK) supplemented with 10% fetal calf serum (Biological Industries Ltd, Cumbernauld, UK), glutamine (Life Technologies Ltd, Paisley, UK), HEPES buffer (Gibco BRL, Paisley, UK), hydrocortisone, insulin, transferrin, and sodium selenite (Sigma Chemical Company Ltd, Poole, UK). HEK293 cells, human embryonic kidney (ATCC^®^ CRL-1573) was maintained in Thermo Scientific HyClone Dulbecco’s Modified Eagle’s Medium (DMEM) supplemented with 10% Thermo Scientific HyClone Defined Fetal Bovine Serum. Cells were grown in an incubator at 37^°^C and an humidified atmosphere of 5% CO_2_. All experiments were performed under serum free conditions to avoid the influence of serum on cell function and investigated events. *L. santarosai* serovar Shermani str. LT821 (ATCC number 43286; pathogenic species) purchased from the American Type Culture Collection (Manassas, VA) was used in this study. These *Leptospira* was propagated at 28^°^C under aerobic conditions in liquid Ellinghausen-McCullough-Johnson-Harris (EMJH) medium (BD Diagnostics) with 10% rabbit serum enriched with L-asparagine, sodium pyruvate, calcium chloride, and magnesium chloride. Bacterial densities were counted with a CASY-Model TT cell counter and analyzer (Roche Innovatis AG; Casy-Technologh, Reutlingen, Germany).

### Construction of TLR2 Over-expressed Plasmid (p-TLR2-Lentip)

The vector containing human *tlr2* gene (pUNO-hTLR2) was obtained from InvivoGen for gene manipulation. The human full-length *tlr2* was amplified by PCR cloning with the primers listed in Table 1 and inserted into the vector pLenti6.3/V5-TOPO to obtain the p-TLR2-Lentip plasmids for stable gene expression in culture cells. PCR product was verified via agarose gel electrophoresis, following PCR purification (Qiagen, Valencia, CA), and will be cloned into the pLenti6.3/V5-TOPO lentiviral vector (Invitrogen). Following TOPO cloning, One Shot Stbl3 Chemically Competent *E. coli* was transformed with DNA. The plasmid DNA with correct size and orientation was verified by restriction enzyme double digestion and confirmed by DNA sequencing.

**Table 1.**
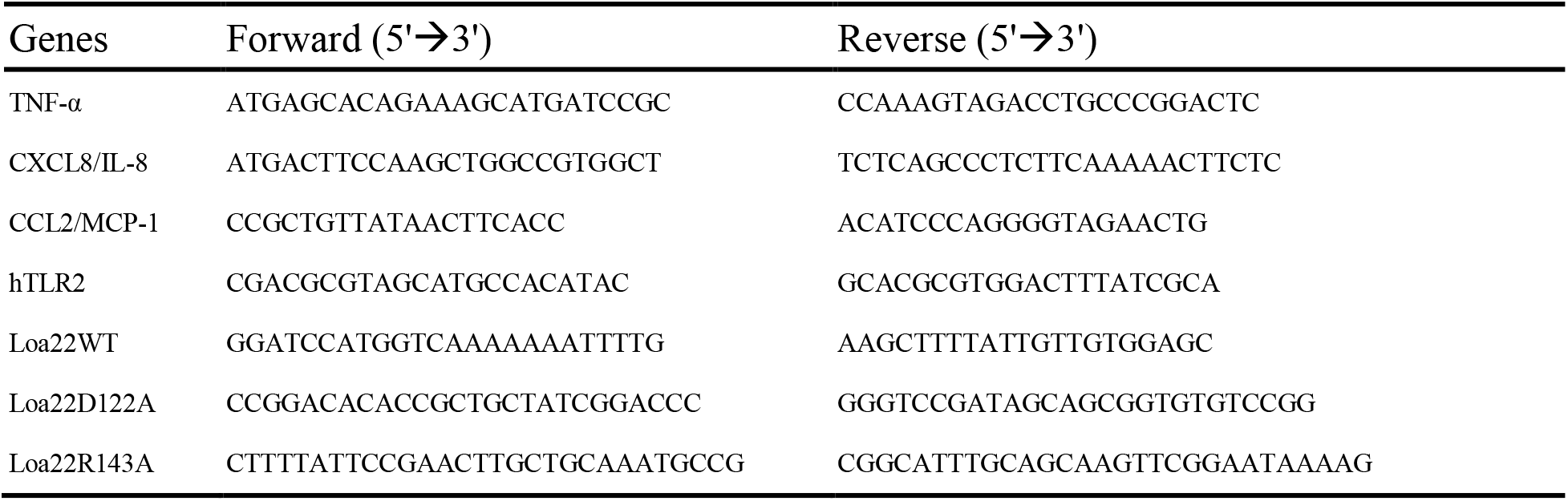
Primers used in this study

### Lentiviral Transduction and Establishment of Stable Cell Line

The p-TLR2-Lentip plasmid was transfected into human embryonic kidney cells, 293FT producer cells with ViraPower packaging mix (pLP1, pLP2, and pLP/vesicular stomatitis virus; ViraPower; Invitrogen) to generate the lentivirus according to manufacturer’s protocol. Human Embryonic Kidney 293 cells (HEK-293 cells; Epithelial Kidney Cells, CRL-1573) were transduced with the lentivirus and stable cell lines were generated by selecting with blasticidin. Cells was grown in DMEM with 10% FBS and maintained in a 37°C humidified atmosphere containing 5% CO_2_. The stable cell line with biological markers (GFP) was confirmed by microscopy-based observation. Cells were collected and named HEK293-TLR2 cells, respectively.

### *Leptospira* Outer Membrane Extraction

Pathogenic *L. santarosai* serovar Shermani and non-pathogenic *L. biflexa* serovar Prato (as a negative control), obtained from the American Type Culture Collection (Manassas, VA), was grown in 10% Ellinghausen McCallough Johnson Harris (EMJH) *Leptospiral* enrichment medium (Detroit, MI). *Leptospira* was cultured for 5–7 days at 28^°^C until they reached a cell density of 10^8^/ml as previously described (5). The *Leptospiral* outer membrane extraction from pathogenic *L. santarosai* serovar Shermani and non-pathogenic *L. biflexa* serovar Prato were extracted with 1% Triton X-114 using the extraction protocol in Haake et al. (33). Briefly, pathogenic and non-pathogenic *Leptospira* was washed in phosphate-buffered saline (PBS) supplemented with 5 mM MgCl_2_ and then extracted at 4^°^C in the presence of 1% protein grade Triton X-114, 10 mM Tris (pH 8.0), 1mM phenylmethylsulfonyl fluoride (PMSF), 1 mM iodoacetamide, and 10 mM ethylenediaminetetraacetic acid. Insoluble material was removed by centrifugation at 17,000 xg for 10 min. Phase separation was conducted by warming the supernatant to 37^°^C and subjecting it to centrifugation for 10 min at 2000 xg. The detergent and aqueous phases were separated and precipitated with acetone and then lyophilized. The extracts was further dissolved in sterile H_2_O, filtered through 0.22 mm membrane filters, and stored at 80^°^C until use.

### Pull-down Assay

HEK293 over-expressed TLR2 cell line (HEK293-TLR2) was used to express the functional full-length human TLR2 protein for pull-down assay. After 48 h cell culture, HEK293-TLR2 cells were incubated with *Leptospira* outer membrane extraction for 4 h and followed by three times PBS washes and centrifugation to remove the non-interacted *Leptospira* outer membrane components. The cells were lysed with cold lysis buffer (1% (w/v) Triton X-100, 150 mM NaCl, 1 mM EDTA, 20 mM Tris-HCl [pH 7.4]) containing protease inhibitor mixture (Roche) for 30 min at 4°C. Lysates were centrifuged for 15 min at 14,000 rpm and supernatants were incubated with mouse anti-V5-tag antibody or mouse IgG (negative control) at 4°C overnight, and then incubated with protein A–agarose beads (Roche) for 4 h. The beads were washed four times with lysis buffer, and the obtained samples were analyzed by 12% (w/v) SDS-PAGE.

### Protein Identification by Mass Spectrometry (MS)

The fractionated pull-down samples (10 μg/lane) from mouse IgG control and anti-V5 antibody were loaded onto 12% (w/v) SDS-PAGE for electrophoresis separation. Protein identification was analyzed by using matrix-assisted laser desorption/ionization time-of-flight (MALDI-TOF) MS, and further validated by MS/MS. Briefly, Protein bands with at least a 2-fold difference were selected for protein identification. Protein bands were manually excised from the SYPRO Ruby – stained preparative SDS-PAGE, washed, and in-gel digested as described before (34). The excised bands were destained using 50 mM NH4HCO3 in 50% (v/v) acetonitrile and dried in a SpeedVac concentrator. Protein was digested by incubation overnight at 37°C with trypsin (5 ng/ ml, Promega, Madison, WI) in 50 mM NH4HCO3 (pH 7.8). Tryptic peptides were extracted from the gel pieces in 1 volume of 0.1% (v/v) TFA while vortexing for 5 min and followed by sonication for 5 min. Protein IDs based on MS and MS/MS data were both searched against the Swiss-Prot database (version 53.0, 252616 protein entries) with species restriction to *Leptospira santarosai* serovar Shermani, by using Mascot 2.1.0 (Matrix Science, London, UK; www.matrixscience.com) to identify the proteins.

### DNA Construction and Mutagenesis

The *loa22* gene (LSS_RS00795; 591bp) was cloned from pathogenic L. santarosai serovar Shermani genomic DNA using pfu-Turbo DNA polymerase (Stratagene, La Jolla, CA, USA). The primers used for *loa22* gene construction were listed in Table 1. After restriction enzyme double digestion, the PCR product was individually inserted into the expression vector pRSET-c (Invitrogen, Groningen, Netherlands). The point mutation of *loa22* variants were obtained by using Q5^^®^^ Site-Directed Mutagenesis Kit (NEB, Ipswich, MA) with their relevant primers. The plasmid DNA was verified by DNA sequencing.

### Bioinformation Analysis

Multiple sequence alignment of Loa22 protein from *L. shermani* and AbOmpA and Pal proteins from *A. baumannii* and *E. coli* were performed with TEXSHADE program (35). The SMART http://smart.embl-heidlbergde/ and LipoP http://www.cbs.dtu.dk/services/LipoP/ web servers were used to search predicted functional and structural domains of Loa22 (16, 36).

### Expression and Purification of Loa22 and Antibody Preparation

The DNA constructs of Loa22 were individually transformed into expression host cell *E.coli* ClearColi^TM^BL21 (DE3) pLys (Lucigen, Middleton, WI). Loa22 and its variants were grown at 37^°^C in the medium with 50 μg/mL ampicillin to an OD_600_ of approximately 0.6. Isopropyl 1-thio-β-galactopyranoside (500 μM) was added with further 4 h grown, *E. coli* cells were harvested by centrifugation at 4,000 *xg* for 15 min and disrupted by sonication in PBS buffer. The cell debris was discarded after centrifugation at 12,000 *xg* for 30 min, and the supernatant was subjected into 9_+_ 9_ļ_ Ni_2+_ -nitrilotriacetic acid (Ni_2+_ -NTA) agarose resin (Qiagen, Valencia, CA, USA) for affinity chromatography purification. The Loa22 and its variants protein were eluted by 250 mM imidazole and stored at -80^°^C for further use. The imidazole was removed by dialysis before assays. rLoa22 antigens were used to induce the production of the polyclonal antibodies (anti-Loa22 antibody) by customized product (Kelowna International Scientific Inc., Taipei, Taiwan).

### RNA Extraction and Real-Time PCR

HEK293/TLR2 were cultured in DMEM/Ham’s F12 medium supplemented with 5% (w/v) fetal calf serum (FCS), 2 mM glutamine, 20 mM HEPES (pH 7.0), 0.4 μg/ml hydrocortisone, 5 μg/ml insulin, 5 μg/ml transferrin, and 28.9 μM sodium selenite at 37^°^C in a humidified atmosphere of 5% (v/v) CO_2_ as previously described (5). Cells were shifted to a serum-free medium for 24 h before adding Loa22 to the cell culture medium. Total RNA was extracted according to the guanidinium thiocyanate/phenol/chloroform method (Cinna/Biotecx Laboratories International Inc., Friendswood, TX) (5, 37). Real-time PCR was executed on the basis of the manufacturer’s instructions using an ABI Prism 7700 with SYBR green I as a double-stranded DNA-specific dye (PE-Applied Biosystems, Cheshire, Great Britain). The primers of *hTNF-α, hCXCL8/IL-8,* and *hCCL2/MCP-1* were shown in Table 1 and constructed to be compatible with a single reverse transcription-PCR thermal profile (95°C for 10 min, 40 cycles at 95°C for 30 s, and 60°C for 1 min). The accumulation of the PCR product was recorded in real time (PE-Applied Biosystems). The results of mRNA levels in different genes are displayed as the transcript levels of the analyzed genes relative to GAPDH (glyceraldehyde-3-phosphate dehydrogenase) transcript level.

### Leptospira PGN Preparation

*Leptospira* PGN was extracted according to the procedure of previous report (38). Briefly, 2 litres of leptospire culture were harvested by centrifugation at 10,000 *xg* for 30 min, washed three times with PBS buffer, and resuspended in 100ml 1% (w/v) SDS in distilled water. The suspension was gently shaken at 37 ^°^C for 18 h and then centrifuged at 110,000 *xg* for 60 min. After a second treatment with 1% (w/v) SDS the pellet was washed three times with 6M urea. The pellet was further resuspended in 100 ml distilled water and collected by centrifugation at 110,000 *xg* to remove the urea. The pellet was suspended in 10 ml 10 mM Tris-HCl buffer (pH 7.4) containing 0.1 mg/ml trypsin and incubated at 37 ^°^C for 18 h. The pellet was collected by centrifugation at 110,000 *xg* for 90 min and the pellet was suspended in 10 ml 10 mM Tris-HCl buffer (pH 7.4) containing 0.1 mg/ml pronase (Sigma) for incubation at 37 ^°^C for 18 h. After digestion at 37 “C for 18 h the pellet was recovered by centrifugation at at 110,000 *xg* for 90 min and then lyophilized. The amount of PGN extracted was measured of the dry weight of pellet and resuspended in distilled water for 1mg/ml and stock at -80 ^°^C until use.

### Enzyme-linked Immunosorbent Assay (ELISA)

The interaction between Loa22 and TLR2 was investigated by ELISA according to the prior method with minor modifications (29, 39). The ELISA plates (Nunc-Immuno Plate; Thermo Scientific, Denmark) were primarily coated with 1 μg TLR2 protein in 100 μl PBS buffer and incubated for 2 h at 37°C. The wells were then washed with PBST (PBS containing 0.05% (v/v) Tween 20) and blocked with 200 μl PBS buffer containing 1% (w/v) BSA for 1 h at 37°C. The plates were incubated overnight at 4°C. Protein samples (Loa22 and its variants; from 0 to 2 μM) in PBS buffer were added to the attached TLR2 protein for 90 min at 37°C followed by a wash with PBST.

The bound Loa22 protein was allowed to interact with 100 μl anti-Loa22 antibody (1:10,000 dilution) for 1 h. The wells were then washed with PBST. Subsequently, 100 μl horseradish peroxidase conjugated donkey anti-rabbit immunoglobulin G (1:5,000 dilution) was loaded and the plates incubated for 1 h at 37°C. After washing with PBST, the 3,3’,5,5’-tetramethylbenzidine (KPL, Gaithersburg, MA) was included for color development for 15 min and then reaction terminated by adding 50 μl 2N H_2_SO_4_. The absorbance was measured at 450 nm in a THERMOmax ELISA reader (Molecular Devices, Sunnyvale, CA). The absorbance at each data point was corrected by subtracting a negative control from BSA coated well. The dissociation constant (K_d_) was calculated according to the prior protocol (40), based on the equation 1:

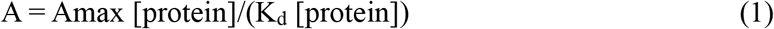

where A is the absorbance at a set protein concentration, Amax is the maximum absorbance for the ELISA plate reader (equilibrium), [protein] is the protein concentration, and Kd is the dissociation equilibrium constant for a given absorbance at a set protein concentration. The binding data were analyzed via SigmaPlot 10.0 program by fitting to the optimal equation (41), assuming that ligands bound to one independent site of a protein interested.

### AFM Tip and Mica Surface Modification

The AFM cantilever made of silicon nitride, PNP-TR (NanoWorld, Neuchâtel, Switzerland), was cleaned by sonication in a series of solvents: 2-propanol, methanol, and deionized water (5 min each), successively. The tips were functionalized according to a previous method with minor modifications (29, 30, 42). At first, the probes were transferred into a 0.1% (v/v) solution of 3-aminopropyltrimethoxysilane (APTES) (Sigma, St. Louis, MO) in toluene for 2 h. Following silanization, the tips were sonicated in methanol and deionized water (5 min each), consecutively. For functionalization, the AFM tips were incubated in a 1% (v/v) solution of glutaraldehyde (Grade II, Sigma) in PBS for 1 h at room temperature. For anchoring proteins, the tips were rinsed with PBS to remove the glutaraldehyde remainders and subsequently incubated with proteins, including wild type Loa22, its variants, and BSA, respectively, for 1 h at room temperature. The functionalized AFM tips were then rinsed three times with PBS to remove the unbound proteins (29, 30). The mica surface was modified for better deposition of proteins interested according to previous report (43, 44). In brief, the mica surface (1 cm^2^) was functionalized with 0.1% (v/v) APTES (Sigma) (AP-mica) followed subsequently by 1% (v/v) glutaraldehyde (Grade II, Sigma) treatment. The proteins (100 ng) were then deposited covalently on the functionalized mica surface for 2 h at room temperature. The unbound proteins were removed and the proteins fixed on the mica were ready for later AFM measurements.

### AFM Force-distance Curves Measurement and Analysis

The functionalized cantilever tips used in this work had a spring constant (k) in the range of 0. 02-0.08 N/m as determined from the amplitude of their thermal vibrations. A commercial atomic force microscope (Nanoscope III, Digital Instruments, Santa Barbara, CA) with a J type scanner was employed throughout this study. The force volume software takes a force curve at each of the 999 points during a two dimensional scan over a sample surface. The X-Y scan size was 150 μm, and the Z scan distance was 5 μm at a rate of 1 Hz. The force applied to the protein modified mica surface was kept below 400 pN. The distance-force curves and force parameters were obtained according to the methods previously described (29, 42). For antibody neutralization analysis, the protein modified mica was treated with anti-TLR2 antibodies (diluted 1:1,000 in PBS) for 1 h followed by five washes with PBS to remove unbound antibodies. The antibody against TLR2 (AbTLR2) was purchased from GeneTex (Irvine, CA). For the binding events, force curves from at least 10 areas on each surface were independently selected for analysis. All the measurements described above were performed with modified tips and showed repeatedly similar results. The force mapping data were transformed into force extension curves for each subsection, and the area under the force extension curvewas then calculated. For dynamic force analysis, the rupture speed was controlled within the range from 14 to 1400 nm/s. SPIP (ImageMetrology A/S,Hørsholm,Denmark)was used for data analysis, and values of extraction force were thus determined with SigmaPlot version 10.0 (SPSS, Chicago, IL). The maximum distribution force was calculated by a Gaussian fit to the force distribution curve. The dynamic profile for maximum distribution force versus the loading rate was obtained from equation

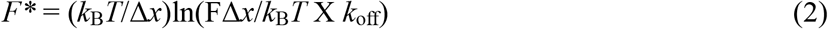

where *F*\* is the most probable rupture force, *k_B_* is the Boltzmann constant, *T* is the temperature, Δ*x* is the potential width of the energy barrier along the direction of the applied force, *k_off_* is the natural dissociation rate at zero force, and F is the loading rate (45).

### Ethics Statement

All animal procedures and experimental protocols were approved by the Institutional Animal Care and Use Committee

## Acknowledgments

The authors thank Microscope Core Laboratory, Chang Gung Memorial Hospital, Linkou, Taiwan.

## Supporting Information Legends

Supplemental Figure S1

AFM image of *L. santarosai* Shermani and force distribution curves of the Loa22 and LPGN. (A) The height model of AFM image for *Leptospira*. The *Leptospira* was spray on mica surface and fixed with 1% Glutaraldehyde for 1 h. The fixed *Leptopsira* was washed with 0.1% TritonX-114 to remove the outer membrane as described in Material and Methods and scanned with AFM in the PBS buffer. (B) 3D view of cell wall structure of *Leptospira.* (C) Force distribution curves of AFM tip interacts to *Leptospira* cell wall, LPGN. (D) Force distribution curves of rLoa22WT modified AFM tip interacts to mica surface. (E) Force distribution curves of rLoa22WT modified AFM tip interacts to *Leptospira* cell wall, LPGN. (F) Force distribution curves of rLoa22D122A modified AFM tip interacts to *Leptospira* cell wall, LPGN. (G) Force distribution curves of rLoa22R143A modified AFM tip interacts to *Leptospira* cell wall, LPGN.

## Author Contributions

Conceived and designed the experiments: SHH CCH CWY. Performed the experiments: SHH CMY YCK LFC. Analyzed the data: SHH CMY YCK CCH CWY. Contributed reagents/materials/analysis tools: SHH CCH YCK LFC. Wrote the paper: SHH CCH CWY.

## Founding Information

This work was supported by the grants from CGMH-NTHU Joint Research CMRPG3E0331 to C.-C.H and CMRPG3H0881 to S.-H.H.; National Science Council, Republic of China: MOST 103-2314-B-182-020-MY3 and MOST 101-2321-B-182-007-MY3 to C.-W.Y. MOST 106-2311-B-182A-001 – and MOST 107-2311-B-182A-003 – to S.-H.H.

